# Unraveling a tangled skein: Evolutionary analysis of the bacterial gibberellin biosynthetic operon

**DOI:** 10.1101/868042

**Authors:** Ryan S. Nett, Huy Nguyen, Raimund Nagel, Ariana Marcassa, Trevor C. Charles, Iddo Friedberg, Reuben J. Peters

## Abstract

Gibberellin (GA) phytohormones are ubiquitous regulators of growth and developmental processes in vascular plants. The convergent evolution of GA production by plant-associated bacteria, including both symbiotic, nitrogen-fixing rhizobia and phytopathogens, suggests that manipulation of GA signaling is a powerful mechanism for microbes to gain an advantage in these interactions. Although homologous operons encode GA biosynthetic enzymes in both rhizobia and phytopathogens, notable genetic heterogeneity and scattered operon distribution in these lineages suggests distinct functions for GA in varied plant-microbe interactions. Therefore, deciphering GA operon evolutionary history could provide crucial evidence for understanding the distinct biological roles for bacterial GA production. To further establish the genetic composition of the GA operon, two operon-associated genes that exhibit limited distribution among rhizobia were biochemically characterized, verifying their roles in GA biosynthesis. Additionally, a maximum-parsimony ancestral gene block reconstruction algorithm was employed to characterize loss, gain, and horizontal gene transfer (HGT) of GA operon genes within alphaproteobacteria rhizobia, which exhibit the most heterogeneity among GA operon-containing bacteria. Collectively, this evolutionary analysis reveals a complex history for HGT of both individual genes and the entire GA operon, and ultimately provides a basis for linking genetic content to bacterial GA functions in diverse plant-microbe interactions.

## INTRODUCTION

The clustering of bacterial biosynthetic genes within operons allows for the controlled co-expression of functionally-related genes under a single promoter, and the opportunity for these genes to be mobilized and co-inherited as a complete metabolic unit via horizontal gene transfer (HGT) [1, 2]. Because operons are responsible for many fundamental biosynthetic pathways in bacteria, analysis of the genetic structure of complex operons can provide important clues regarding the selective pressures driving the evolution of bacterial metabolism, and can also give insight into the occurrences and mechanisms of HGT.

The ability for bacteria to produce gibberellin (GA), a ubiquitous plant hormone, is imparted by a GA biosynthetic operon (GA operon; **Figure 1**), which is found in both nitrogen-fixing rhizobia and phytopathogenic bacteria [3–5]. While the diterpenoid GA phytohormones act as endogenous signaling molecules for growth and development in vascular plants [6], plant-associated fungi and bacteria have convergently evolved the ability to produce GA as a mechanism for host manipulation [4, 7–9]. The phenomenon of GA production by plant-associated microbes has important biological implications, as perturbation in GA signaling can lead to extreme phenotypic changes in plants. For example, production of GA by the rice pathogen *Gibberella fujikuroi* leads to dramatic elongation and eventual lodging of rice crops [10], and impaired GA metabolism is responsible for the semi-dwarf crop phenotypes associated with crops utilized within the Green Revolution [11, 12]. More recently, it has been shown that GA acts as a virulence factor for phytopathogenic bacteria [9], and can affect nodulation phenotypes when produced by rhizobia in symbiosis with legumes [4]. Therefore, studying the biosynthesis and biological function of microbial GA is crucial to our understanding of how these plant-microbe interactions can affect plant health and development.

**Figure 1.**
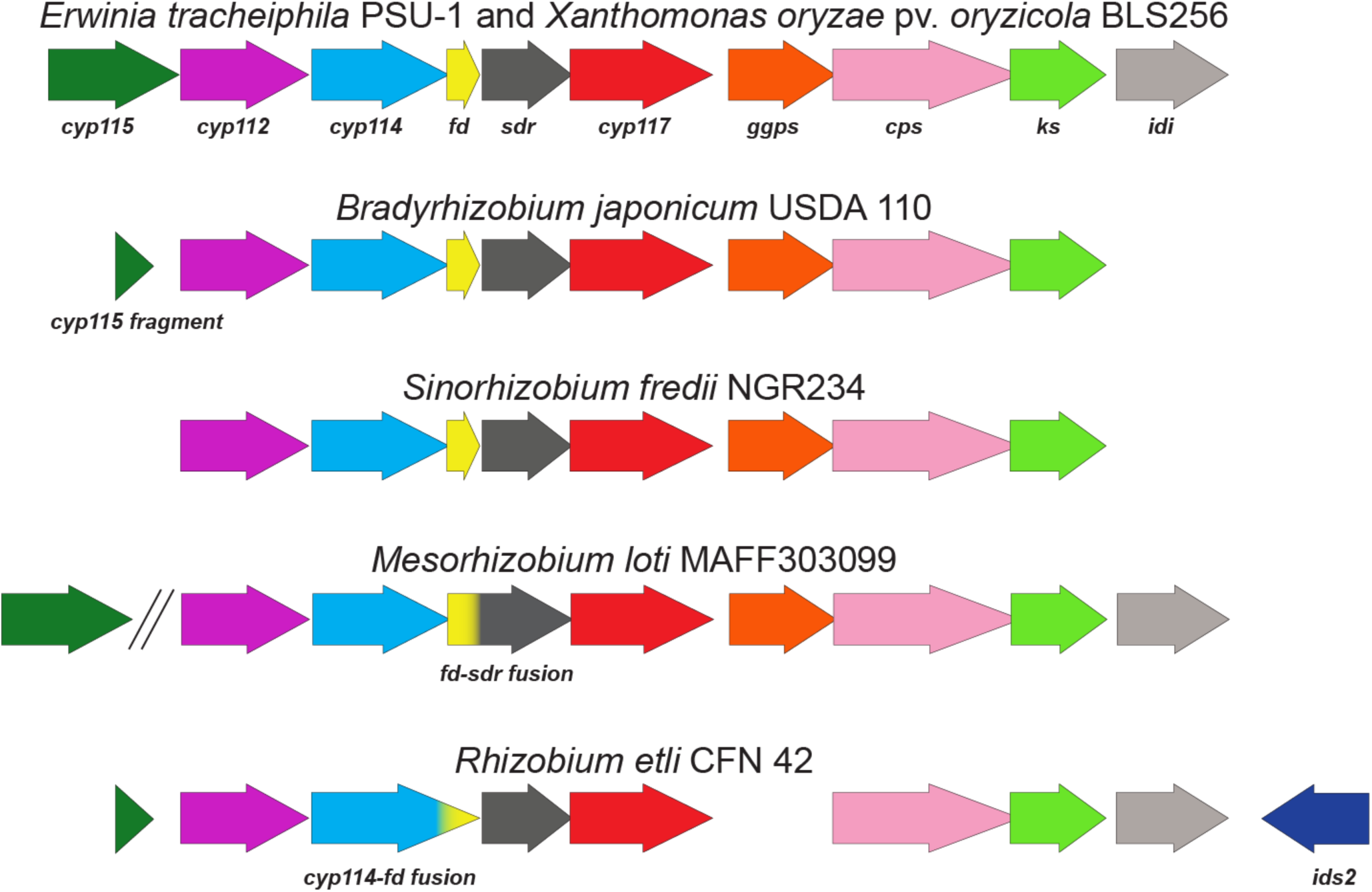
Diversity among GA biosynthetic operons in divergent bacterial lineages. The core operon genes are defined as *cyp112, cyp114, fd, sdr, cyp117, ggps, cps*, and *ks*, as these are almost always present within the GA operon. Other genes, including *cyp115, idi*, and *ids2*, exhibit a more limited distribution among GA operon-containing species. The tandem diagonal lines in the *Mesorhizobium loti* MAFF303099 operon indicates that *cyp115* is not located adjacent to the rest of the operon.

The GA operon was discovered in the rhizobial symbiont of soybean, *Bradyrhizobium diazoefficiens* USDA 110 [13], This operon contains a geranylgeranyl diphosphate synthase (*ggps*), two diterpene synthases/cyclases (*cps* and *ks*), three cytochrome P450 (CYP) monooxygenases (*cyp112, cyp114*, and *cyp117*), a short-chain dehydrogenase/reductase (*sdr*_*GA*_), and a ferredoxin (*fd*_*GA*_) [13, 14]. The *B. diazoefficiens* operon also contains a severely truncated, presumably non-functional CYP gene (pseudo *cyp115*, or *p-cyp115*) located at the 5’ end of the operon. The core gene cluster, which contains all of the aforementioned genes other than *cyp115*, is widely distributed in symbiotic nitrogen-fixing rhizobia from the alphaproteobacteria class (α-rhizobia) [15], and biochemical characterization of GA operon genes in several α-rhizobia, including *B. diazoefficiens, Sinorhizobium fredii*, and *Mesorhizobium loti*, has demonstrated that this core operon is responsible for biosynthesis of GA_9_, the penultimate intermediate to the bioactive phytohormone GA_4_ [3, 4, 16–18]. While exclusively found in plant-associated bacteria [19], the GA operon exhibits scattered distribution within the α-rhizobia, and functional versions of the operon can also be found in several betaproteobacterial rhizobia symbionts (β-rhizobia) [20, 21]. Analogous GA operons can be found in certain gammaproteobacterial plant pathogens as well (e.g. *Xanthomonas* and *Erwinia* species), and characterization of the GA operon from several distant gammaproteobacterial lineages has demonstrated that the biosynthetic functionality of this operon is conserved [5, 9, 20].

The abundance of sequenced bacterial genomes indicates that the GA operon structure is more complex and variable than that initially described for the α-rhizobia *B. diazoefficiens* in which this operon was initially identified [14, 15]. Specifically, certain bacteria with the GA operon were found to contain a full length *cyp115* gene at the 5’ end of the gene cluster, as opposed to a pseudo-gene/fragment, and this enzyme has been shown to catalyze the final step in bioactive GA biosynthesis, converting GA_9_ into bioactive GA_4_ [5, 20, 22]. Additionally, many bacterial strains possess a putative isopentenyl diphosphate δ-isomerase (*idi*) gene located at the 3’ end of the operon, which presumably functions in balancing the concentrations of the (di)terpenoid building blocks, isopentenyl diphosphate (IPP) and dimethylallyl diphosphate (DMAPP) [23]. Full-length *cyp115* and *idi* genes are notably absent from many α- and β-rhizobia with the operon, while copies of these genes are essentially always present in the GA operons of gammaproteobacterial phytopathogens (**Figure 1**). Intriguingly, it appears that some of the α-rhizobia have specifically lost these genes, as fragments of both *cyp115* and *idi* can be found flanking the core gene cluster in many of the relevant species/strains [22, 24]. Moreover, a small number of α-rhizobia have a presumably inactivating frameshift mutation in the canonical *ggps* within their operon, but have an additional isoprenyl diphosphate synthase (IDS) gene adjacent to the operon (*ids2*) [17], which could potentially compensate for the loss of *ggps*. Overall, this heterogeneity of the GA operon in rhizobia provides an excellent opportunity for analyzing the formation and reorganization of bacterial gene clusters.

Initial phylogenetic analyses of the GA operon suggested that it may have undergone HGT among bacterial lineages [17, 20]. Furthermore, the varying genetic structure of the operon in divergent species, including both symbionts and pathogens, suggests that selective pressures unique to certain bacteria may be driving the acquisition or loss of not only the GA operon, but also some of the associated genes. Thus, detailed analysis of GA operon evolution will help elucidate the evolutionary processes that have shaped bacterial GA biosynthesis in plant-microbe interactions. Here, the predicted biochemical functions were assessed and confirmed for the *idi* and *ids2* genes that are sporadically associated with the GA operon, thereby providing evidence for their roles in GA biosynthesis. This clarification of genetic content prompted further analysis of the distribution and function of the GA operon in bacteria more generally, thereby providing an overview of the genetic diversity and evolutionary history of this gene cluster. Using an algorithm developed to analyze the assembly and evolution of gene blocks (i.e. genes within operons/clusters) [25], the distribution and phylogeny of the GA operon was further analyzed within the α- rhizobia, which display a large amount of diversity in operon structure and genetic content. Altogether, this thorough assessment of the underlying genetics and biochemistry of the GA operon allows for the formulation of informed hypotheses regarding the biological function of GA production within diverse bacterial lineages.

## RESULTS

### Biochemical characterization of two accessory GA operon genes

Given that most bacteria do not normally produce (*E,E,E*)-geranylgeranyl diphosphate (GGPP), a necessary precursor, GA biosynthesis requires the presence of a *ggps* gene. In the GA operon-containing strain *Rhizobium etli* CFN 42, the operon *ggps* contains a frameshift mutation that results in a severely truncated protein [17]. However, a second predicted IDS gene *(ids2)*, albeit with low sequence identity to the canonical operon *ggps* found in other *Rhizobium* species (<30% at the amino acid level), is found in close proximity to this strain’s operon (**Figure 1**), as well as in several other α-rhizobia in which *ggps* similarly appears to be inactive. Given the conservation of these modified operons, we hypothesized that the encoded IDS2 also produces GGPP, thereby restoring functionality to these GA operons. Indeed, recombinantly expressed IDS2 from *R. etli* CFN42 (*Re*IDS2) produced GGPP as its sole product from the universal isoprenoid precursors IPP and DMAPP (**Supplementary Figure 1**). Thus, IDS2 can functionally complement the loss of the canonical *ggps* to restore production of GA in these operons. Accordingly, hereafter these *ids2* gene orthologs are referred to as *ggps2* to reflect their biochemical function (e.g. *Re*IDS2 becomes *Re*GGPS2).

The only remaining gene strongly associated with the GA operon but not yet characterized was *idi*, which has been presumed to be involved in balancing the ratio of IPP and DMAPP isoprenoid building blocks for diterpenoid biosynthesis [23]. The GA operon *idi* from *Erwinia tracheiphila* (*Et*IDI), a gammaproteobacteria plant pathogen, was cloned and heterologously expressed in *E. coli*. To test for activity, a coupled enzyme assay with *Et*IDI and *Re*GGPS2 was employed. Because IDS enzymes require both IPP and DMAPP as substrates, *Re*GGPS2 is unable to produce GGPP with only IPP or DMAPP alone as substrate. Addition of *Et*IDI into these reactions enabled the production of GGPP by *Re*GGPS2 from either IPP or DMAPP alone (**Supplementary Figure 2**), thus indicating that *Et*IDI can effectively interconvert these.

### HGT of the GA operon within alphaproteobacterial rhizobia

The scattered distribution of the GA operon among three classes of proteobacteria suggests HGT of this gene cluster. Previous phylogenetic analysis suggests that the ancestral gene cluster initially evolved within gammaproteobacterial phytopathogens, as their operon genes exhibit greater phylogenetic divergence than those in the rhizobia, and that the operon was subsequently acquired by α- and β- rhizobia in separate HGT events [20]. Additionally, specific phylogenetic analysis of the GA operon within α- rhizobia suggests that it may have subsequently undergone additional HGT within this class [17].

It has previously been noted that the GC content of the GA operon in rhizobia is particularly high compared with the surrounding genomic sequence [14, 24, 26, 27], a phenomenon that is often associated with HGT [28]. To better assess the increased GC content of the GA operon, we analyzed the gene cluster sequences and the surrounding DNA in exemplaries from four of the major α-rhizobia genera (*Bradyrhizobium, Mesorhizobium, Rhizobium*, and *Sinorhizobium*) and two genera of the gammaproteobacterial plant pathogens (*Erwinia* and *Xanthomonas*). In each case, the GA operon has noticeably higher GC content than the surrounding DNA (>10% higher), with sharp drops in GC content preceding and following the operon (**Supplementary Figure 3**). Further support for HGT of the GA operon has been suggested by the presence of insertional sequence (IS) elements flanking the operon (e.g. transposases and integrases) in many species [5, 22]. Overall, these collective observations strongly support HGT of the GA operon, consistent with its widely-scattered distribution throughout the proteobacteria.

As an added layer of complexity, it is generally accepted that the large symbiotic or pathogenic genomic islands or plasmids (i.e. symbiotic or pathogenic modules), which are associated with the plant-associated lifestyle of the bacteria in question, are capable of undergoing HGT [29]. For α-rhizobia strains where sufficient genomic information is available, the GA operon is invariably found within the symbiotic module [24, 26, 27, 30–32]. Thus, there may be multiple levels of HGT with the GA operon in α-rhizobia: one in which the entire symbiotic module, including a GA operon, is transferred, and one in which the GA operon alone is transferred. Indeed, this double-layered HGT for the GA operon has been previously suggested based on phylogenetic incongruences between genes representative of species (16S rRNA), symbiotic modules (*nifK*), and GA operon (*cps*) similarity [17].

### Gene cluster analysis

While the GA operons found in gammaproteobacteria exhibit essentially uniform gene content and structural composition, those from the α-rhizobia exhibit much more diversity in genetic structure. This suggests that selective pressures specific to the rhizobia, presumably their symbiotic relationship with legumes, may have driven this heterogeneity in the operon. To better understand the evolutionary history of the GA operon in the α-rhizobia, a more thorough analysis was carried out with Reconstruction of Ancestral Genomes Using Events (ROAGUE) software [25, 33]. ROAGUE generates a phylogenetic tree with selected taxa that contain gene blocks (i.e. gene clusters) of interest, and then uses a maximum parsimony approach to reconstruct a predicted gene block structure at each ancestral node of the tree. Using the ROAGUE method, the evolutionary events involved in the genetic construction of orthologous GA operon gene blocks in the α-rhizobia, specifically gene loss, gain, and duplication, were quantitatively assessed (see **Supplementary Figure 4** for a summary of the method pipeline). A total of 118 α-rhizobia with the GA operon were included in this analysis. The most phylogenetically distant GA operon to those in the α-rhizobia is found within *E. tracheiphila*, and as such this was used as an outgroup. Additionally, to observe the relative relationship between alpha- and gamma-proteobacterial operons, the GA operon from *Xanthomonas oryzae* was also included in the analysis.

An initial reconstruction was made by creating a species tree using the amino acid sequence of *rpoB* (RNA polymerase β subunit) from each strain as the phylogenetic marker gene (“full species tree” or *FS*) (**Supplementary Figure 5**). However, the species tree is rarely indicative of a given gene’s evolution, and even less so concerning operon evolution where HGT is involved. To better understand the evolution of the GA operon in relationship to the bacterial species, we constructed a second tree with concatenated protein sequences comprising the core GA operon (“full operon tree” or *FO*) (**Supplementary Figure 6**). Due to the large number of species being analyzed, along with apparent phylogenetic redundancy that could introduce bias, reconstructions were also made with only the more distinct representative strains by using the Phylogenetic Diversity Analyzer (PDA) program, which reduced the number of analyzed taxa to 64 [34]. These reduced phylogenetic trees are referred to as the “partial species tree” or *PS* (**Figure 2**), and the “partial operon tree” or *PO* (**Figure 3**).

**Figure 2.**
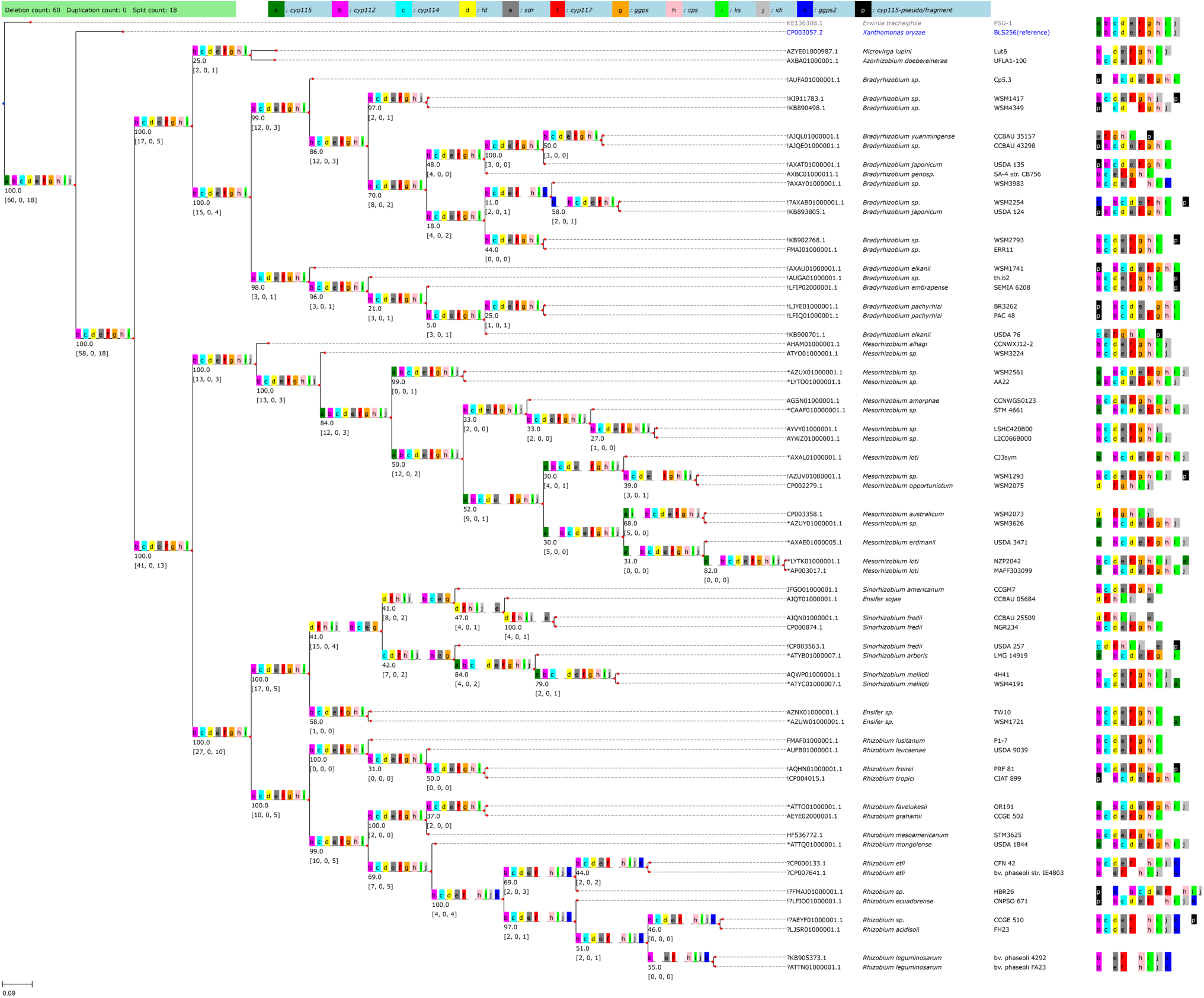
Reduced ancestral reconstruction of the GA biosynthetic operon using *rpoB* for phylogenetic analysis. A phylogenetic tree was constructed with alignments of *rpoB* protein sequences from 118 α-rhizobia species and two gammaproteobacteria using the Neighbor-Joining method as a measure of distance between species. The number of analyzed species was reduced to 64 with the Phylogenetic Diversity Analyzer software, and ROAGUE was then applied to create the ancestral operon reconstruction. The lower-case letters in each tree node represent the genes in the orthoblock (e.g. “a” represents “*cyp115*”), with each gene additionally indicated by a unique color (see legend at top of figure). A blank space between genes designates a split ≥500bp between the genes to either side of the blank space. The green bar on the top left of the figure displays the total number of events that occur in this reconstruction. For each inner node *u*, the floating number (e.g. 98.0) represents the bootstrap value of the tree. The numbers in the brackets indicate the cumulative count of events going from the leaf nodes to node *u* in the following order: [deletions, duplications, splits]. Each leaf node is accompanied with symbols (*, ?, !), the genomic accession number, the species/strain name, and the gene block for that strain. An asterisk (*) indicates the gene block contains full length *cyp115* (gene “a”); an exclamation (!) indicates that the gene block contains a truncation/fragment of *cyp115*, and a question mark (?) indicates the gene block contains *ggps2* (gene “k”). The reference strain, *Xanthomonas oryzae* pv. *oryzicola* BLS256, is in blue, and the outgroup strain, *Erwinia tracheiphila* PSU-1, is in gray. These naming and color conventions persist through this study.

**Figure 3.**
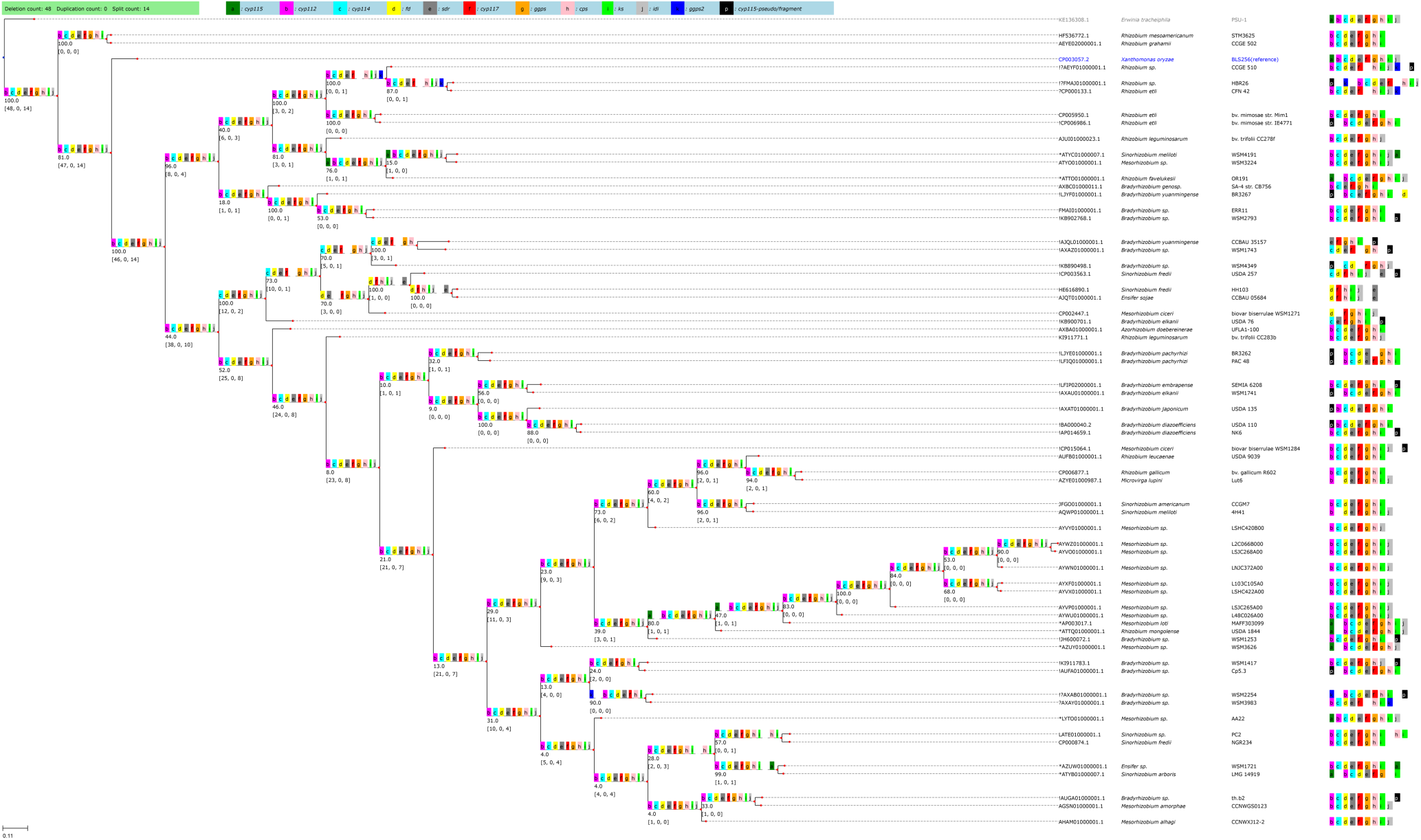
Reduced ancestral reconstruction of the GA biosynthetic operon using the concatenated operon for phylogenetic analysis. A phylogenetic tree was constructed with alignments of concatenated proteins from core GA operon genes (*cyp112-cyp114-fd-sdr-cyp117-cps-ks*) from 118 α-rhizobia species and two gammaproteobacteria using the Neighbor Joining method. The number of analyzed species was reduced to 64 with the Phylogenetic Diversity Analyzer software, and ROAGUE was then applied to create the ancestral operon reconstruction. and ROAGUE was then applied to create the ancestral operon reconstruction. All annotations are described previously in **Figure 2**.

The ability of different ancestral reconstructions to capture the likely vertical evolution of a gene cluster can be assessed by the number of events (loss, gain, and duplication) calculated by this method, with a lower number of events indicating a more parsimonious reconstruction. From this analysis, it was found that fewer evolutionary events are reconstructed in *FO* (75 events) than in *FS* (121 events) (**Supplementary Figures 5 & 6**), with the same relative trend observed with the partial trees (62 events for *PO* vs. 78 events for *PS*) (**Figures 2 & 3**). The greater parsimony (i.e. fewer reconstructed events) observed in reconstructions built with alignments of the concatenated GA operon strongly supports the previously suggested hypothesis of HGT among α-rhizobia [17]. Accordingly, the reconstructions based on GA operon similarity (i.e. *FO* and *PO*) were used for further analyses of operon inheritance.

In contrast to the phytopathogens, a full-length *cyp115* gene is absent from the genomes of most rhizobia (including both α- and β- rhizobia), which typically have only the core operon and thus can only produce the penultimate intermediate GA_9_ rather than bioactive GA_4_ [18, 20, 22]. ROAGUE analysis indicates that *cyp115* loss occurred soon after α-rhizobia acquisition of the GA operon, as the reconstructed ancestral node that connects the α-rhizobia to *X. oryzae* (and the rest of the gammaproteobacteria) does not contain *cyp115* (**Figure 3**). Although the α-rhizobia presumably acquired their GA operon from a gammaproteobacterial ancestor, the gammaproteobacteria seem to always have *cyp115* at the 5’ end of the operon. In contrast, the α-rhizobia typically only have a partial *cyp115* pseudo-gene/fragment located at this position, as previously described [14, 22]. This suggests that the original operon acquired by an α- rhizobia ancestor contained *cyp115*, and that this gene was subsequently lost.

Although most rhizobia have lost *cyp115*, a subset of α-rhizobia (<20%) with the GA operon also have a full-length, functional *cyp115*. However, only in one strain (*Mesorhizobium* sp. AA22) does the GA operon have *cyp115* in the same location as in gammaproteobacterial GA operons [22]. Strikingly, ROAGUE analysis indicates that full-length *cyp115* has been regained independently in at least three different lineages, which is apparent in either the *PS* or *PO* reconstructions (**Figures 2 & 3**). Indeed, other than in *Mesorhizobium* sp. AA22, these full-length *cyp115* reside in alternative locations relative to the rest of the GA operon (e.g. 3’ end of operon, or distally located), as previously described [22], which further supports independent acquisition via an additional HGT event.

Unlike *cyp115*, the *idi* gene seems to have been more widely retained by α-rhizobia, as >50 of these strains possess this gene, which seems to invariably exhibit analogous positioning – i.e. as found in the gammaproteobacterial GA operons. This indicates loss of *idi* in many strains, albeit with notable differences among the major α-rhizobia genera. For example, while the presence of *idi* appears to be almost random within *Rhizobium* (16/26 strains) and *Sinorhizobium*/*Ensifer* (8/14), it is nearly absent from all *Bradyrhizobium* (2/40), but ubiquitously found in *Mesorhizobium* (36/36).

Not surprisingly, *ggps2* seems to be invariably associated with operons in which the canonical *ggps* is inactive (**Figures 2 & 3**), and is only found in 13 α-rhizobia (of the 118 strains analyzed here). However, the ancestral reconstructions further indicate that *ggps2* is present in at least two distinct clades in all trees; one composed of closely related *Rhizobium* strains, and another with two *Bradyrhizobium* strains. While the *Rhizobium* all have homologous mutations in *ggps*, with similar positioning of *ggps2* (within 500 bp of the 3’ end of the operon), the two *Bradyrhizobium* have distinct *ggps* mutations, with *ggps2* positioned on opposite sides of the operon. This suggests that, following initial acquisition of *ggps2*, this was further propagated via additional HGT events, in each case to complement inactivation of the canonical *ggps*, along with subsequent vertical transmission at least in *Rhizobium*, similar to the observed re-acquisition of *cyp115* noted above.

In addition to ancestral gene loss and gain events, there further have been fusions between neighboring biosynthetic genes within the GA operon. In some α-rhizobia, the *fd*_*GA*_ gene, which is usually a distinct coding sequence, is found in-frame with either the 5’ proximal *cyp114* gene, or the 3’ proximal *sdr*_*GA*_ gene, resulting in either *cyp114*-*fd* or *fd-sdr* fusions, which presumably encode bifunctional proteins. As fusion events are not analyzed by ROAGUE, these were assessed and categorized manually (**Supplementary Tables 1 & 2**). The *cyp114-fd* fusion is only found in a single clade consisting almost entirely of *Rhizobium* species, which is most evident in the *FO* reconstruction (**Supplementary Figure 6**). By contrast, while the *fd-sdr* fusion is largely found in a clade consisting of mostly *Mesorhizobium* species (**Supplementary Figure 6**), including *M. loti* MAFF303099 where activity of the fused Fd-SDR has been biochemically verified [4], such fusions appear to have independently occurred in other clades of α- rhizobia. Beyond these multiple observations in α-rhizobia, it should be noted that a *fd-sdr* fusion appears to have independently arisen in the β-rhizobia as well [21], further indicating that this is not functionally problematic.

## DISCUSSION

Collectively, our analyses demonstrate a complex history of GA operon function, distribution, and evolution within the proteobacteria (**Figure 4**). Critical to this analysis was characterization of the *ggps2* and *idi* genes. Although these were previously noted to be associated with the GA operon, their function had not yet been demonstrated. To our knowledge, the *ggps2* and *idi* genes were the only remaining uncharacterized genes associated with the GA operon, and thus characterizing the enzymes encoded by these genes represents the final step in elucidation of the associated biosynthetic capacity. Given that bacteria typically produce both isoprenoid precursors IPP and DMAPP directly via the methyl-erythritol-phosphate (MEP) pathway [23], an IDI is not strictly required, though it is possible that the presence of the *idi* gene would allow for increased flux towards GA by balancing precursor supply. Since the *idi* gene is ubiquitous in phytopathogen GA operons and has been lost multiple times in rhizobia, it may be that this gene optimizes GA production, which presumably assists use of GA as a virulence factor by the phytopathogens. However, the utility of optimized GA production by rhizobia is not evident, and thus it is not clear why some α-rhizobia lineages retain this gene.

**Figure 4.**
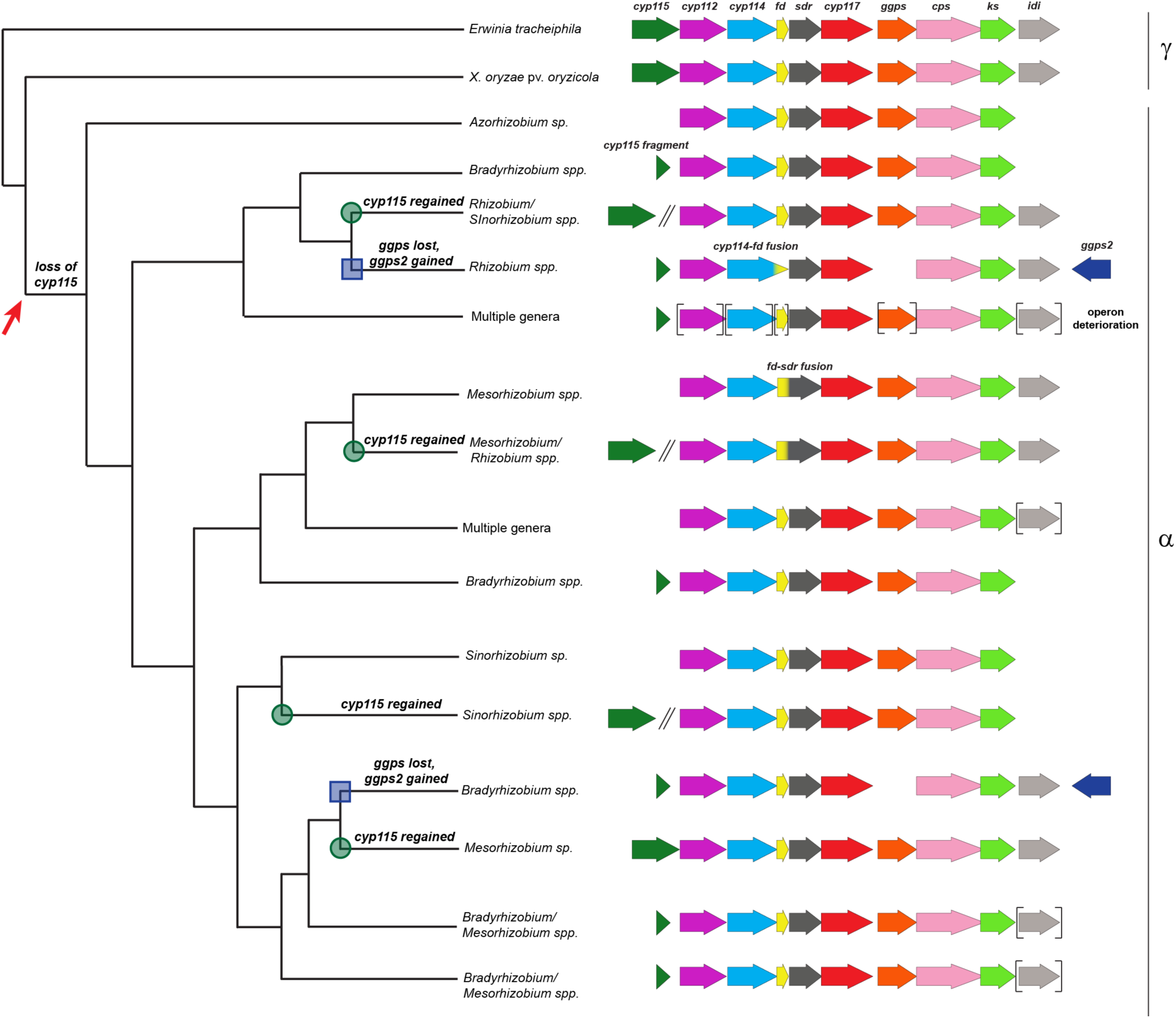
Summary of ancestral reconstruction for the GA biosynthetic operon. As a representation of GA operon evolution, the results from the full reconstruction generated using the concatenated operon (*FO*) are summarized here. Initial loss of the *cyp115* gene is indicated with a red arrow, while reacquisition of this gene is indicated with a green circle at the ancestral node. Loss of *ggps* and acquisition of *ggps2* is indicated by a blue box at the ancestral node. Brackets around a gene represent variable presence within that lineage. Double slanted lines indicate genes that are not located within the cluster (i.e. >500 bp away). The family of proteobacterial lineages is indicated to the right of the figure (α and γ labels).

Unlike the isoprenoid precursor molecules, GGPP is not normally produced by most bacteria, and thus verification of *ggps2* as a GGPP synthase clarifies that GA biosynthesis is still possible in rhizobia where the original operon *ggps* is no longer functional. Interestingly, the *ggps2*-containing *Rhizobium* lineage also harbors a previously defined mutation in the *cps* gene that has been shown to affect product outcome [35]. In particular, the otherwise conserved asparagine from the catalytic base dyad is replaced with a serine in this lineage, which results in predominant production of a distinct compound unrelated to GA biosynthesis (8β-hydroxy-*ent*-copalyl diphosphate), along with small amounts of the relevant GA intermediate (*ent*-copalyl diphosphate). Although retention of the operon indicates that the associated production of GA still provides a selective advantage to these *Rhizobium* strains, despite the presumably reduced flux, it is tempting to speculate that this observation reflects genetic drift of the *cps* in the interlude between loss of *ggps* and acquisition of *ggps2*.

The ROAGUE analysis reported here is consistent with the hypothesis that the GA operon has undergone HGT between various plant-associated bacteria, including phytopathogenic gammaproteobacteria and symbiotic, nitrogen-fixing α- and β- rhizobia. Indeed, there appear to be three layers of HGT relevant to GA production that occur within the α-rhizobia: 1) acquisition of the symbiotic module (i.e. symbiotic plasmid or genomic island), either with or without the GA operon, the latter of which can be followed by 2) separate acquisition of the GA operon within the symbiotic module, with the GA operon enabling 3) subsequent acquisition of auxiliary genes, including *ggps2* and, more interestingly, *cyp115*. Although widespread within proteobacteria, the GA operon has thus far only been found in plant-associated species [19]. While this is not surprising due to the function of GA as a phytohormone, it emphasizes that such manipulation of host plants is an effective mechanism for bacteria to gain a selective advantage. Indeed, the ability to produce GA seems to be a powerful method of host manipulation for plant-associated microbes more generally, as certain phytopathogenic fungi also have convergently evolved the ability to produce GA as a virulence factor [8, 36].

Despite wide-ranging HGT of the GA operon between disparate classes of proteobacteria, its scattered distribution within each of these classes strongly indicates that the ability to produce GA only provides a selective advantage under certain conditions. This is evident for both symbiotic rhizobia and bacterial phytopathogens. For example, the GA operon is selectively found in the *oryzicola* pathovar of *X. oryzae*, where the resulting GA acts as a virulence factor suppressing the plant jasmonic acid (JA) induced defense response [9, 37, 38]. By contrast, production of GA by the α-rhizobia *M. loti* MAFF303099 limits the formation of additional nodules, apparently without a negative impact on plant growth [4].

The occurrence of GA operon fragments (i.e. presence of some, but not all necessary biosynthetic operon genes) in many rhizobia indicates that production of GA is not advantageous in all rhizobia-legume symbioses. For example, at the onset of this study we identified >160 α-rhizobia with an obvious homolog of at least one GA operon gene, yet only ∼120 of these contained a gene cluster (i.e. two or more biosynthetic genes clustered together), and ∼20% of these clusters (26 of the 120 α-rhizobia operons analyzed here) are clearly non-functional due to the absence of key biosynthetic genes, consistent with dynamic selective pressure. It has been suggested that the GA operon is associated with species that inhabit determinate nodules [17], as these nodules grow via cell expansion (an activity commonly associated with GA signaling [39]), rather than indeterminate nodules, which grow via continuous cell division [40]. However, while the presence of the GA operon does seem to be somewhat enriched within rhizobia that associate with determinate nodule-forming legumes, there are many examples of rhizobia with complete GA operons that were isolated from indeterminate nodules. For example, while most GA operon-containing *Bradyrhizobium* species associate with determinate nodule-forming plants, many species from the *Ensifer*/*Sinorhizobium, Mesorhizobium*, and *Rhizobium* genera with the operon were isolated from indeterminate nodules, as were all three of the β-rhizobia with the GA operon (Integrated Microbe Genomes, JGI). This raises the question of why only some rhizobia have acquired and maintained the GA operon, and thus the capacity to produce GA.

In addition to its scattered distribution, the operon exhibits notable genetic diversity within the α- rhizobia. For example, ROAGUE analysis indicates that loss of the usual *ggps* and subsequent recruitment of *ggps2* has been followed by HGT of this to other operons in which *ggps* has been inactivated, while *idi* appears to have independently lost several times. Similarly, fusion of *fd*GA with either the preceding *cyp114* or following *sdr*GA also appears to have occurred multiple times in α-rhizobia, and certainly separately in β-rhizobia [21]. Although loss of *idi* and such gene fusions may affect the rate of GA production, it appears that this can be accommodated in the symbiotic rhizobia-legume interaction. Indeed, the expression of the GA operon is delayed in this relationship [18], perhaps to mitigate any deleterious effects of GA during early nodule formation, which has been shown to be inhibitory to nodule formation, at least at higher concentrations [41].

Perhaps the most striking evolutionary aspect of the rhizobial GA operons is the early loss and scattered re-acquisition of *cyp115* in α-rhizobia. While almost all α-rhizobia GA operons contain only remnants of *cyp115* at the position in the GA operon where it is found in gammaproteobacteria phytopathogens [22], there is one strain (*Mesorhizobium* sp. AA22) where a full-length copy is found at this location. Phylogenetic analysis further suggests that this *cyp115* from *Mesorhizobium* sp. AA22 is closest to the ancestor of all the full-length copies found in α-rhizobia, which are otherwise found at varied locations relative to the GA operon [22]. The ROAGUE analysis reported here indicates that *cyp115* was lost shortly after acquisition of the ancestral GA operon by α-rhizobia, despite full-length copies being present in several different lineages. Accordingly, these results support the hypothesis that *cyp115* has been re-acquired by this subset of rhizobia via independent HGT events. Notably, while not recognized in the original report [4], this includes *M. loti* MAFF303099, the only strain in which the biological role of rhizobial production of GA has been examined. Because *cyp115* is required for endogenous production of bioactive GA_4_ from the penultimate (inactive) precursor GA_9_, this highlights the question of the selective pressures driving evolution of GA biosynthesis in rhizobia.

The contrast between GA operon-containing bacterial lineages provides a captivating rationale for the further scattered distribution of *cyp115* in rhizobia. In particular, the phytopathogens all contain *cyp115* and are thus capable of direct production of bioactive GA_4_, which serves to suppress the JA-induced plant defense response [9]. This observation naturally leads to the hypothesis that rhizobial production of GA_4_ might negatively impact the ability of the host plant to defend against microbial pathogens invading the roots or root nodules, which would compromise the efficacy of this symbiotic interaction. Such detrimental effect of rhizobial production of bioactive GA_4_ may have driven loss of *cyp115*. However, this would also result in a loss of GA signaling, as GA_9_, the product of an operon missing *cyp115*, presumably does not exert hormonal activity [42]. One possible mechanism to compensate for *cyp115* loss would be legume host expression of the functionally-equivalent plant GA 3-oxidase (GA3ox) gene (from endogenous plant GA metabolism) within the nodules in which the rhizobia reside. Expression of this plant gene would alleviate the necessity for rhizobial symbiont maintenance of *cyp115*, and would further allow the host to control the production of bioactive GA_4_, and thereby retain the ability mount an effective defense response when necessary. Re-acquisition of *cyp115* might then be driven by a lack of such GA3ox expression in nodules by certain legumes. However, this scenario remains hypothetical - though precisely controlled GA production by the plant has been shown to be critical for normal nodulation to occur [41, 43], coordinated biosynthesis of GA_4_ by rhizobia and the legume host would need to be demonstrated. This includes both the transport of GA_9_ from microbe to host plant, as well as subsequent conversion of this precursor to a bioactive GA (e.g. GA_4_). Accordingly, continued study of the GA operon will provide insight into the various roles played by bacterially-produced GA in both symbiotic rhizobia-legume relationships, as well as antagonistic plant-pathogen interactions, which in turn can be expected to provide fundamental knowledge regarding the ever-expanding roles of GA signaling in plants.

## METHODS

### Biochemical characterization of *Re*IDS2 and *Et*IDI

*Re*IDS2 and *Et*IDI were cloned from *Rhizobium etli* CE3 (a streptomycin-resistant derivative of *R. etli* CFN42) [44] or *Erwinia tracheiphila* PSU-1, respectively, into pET101/D-TOPO (Invitrogen). The resulting 6xHis-tagged expression constructs were utilized to generate recombinant enzymes that were purified via Ni-NTA agarose (Qiagen). IDS enzyme assays were carried out in triplicate as previously described [45]. Detailed protocols for these experimental procedures can be found in the Supplemental Information document.

### Operon phylogenetic reconstruction

#### Data acquisition

Initial BLAST analysis (on April 12, 2017) revealed 166 bacterial strains that contain homologs of one or more of the GA operon genes. Given a set of 166 species/strain names, the corresponding genome assembly files were retrieved from the NCBI website. Using their assembly_summary.txt file, the strains’ genomic fna (fasta nucleic acid) files were downloaded. The number of strains analyzed was further reduced by only including strains with multiple GA operon genes (>2) clustered together, resulting in a final total of 118 strains. Retrieved genome assemblies for these strains were then annotated using Prokka [46].

#### Identifying orthologous gene blocks

The terms reference taxa, neighboring genes, gene blocks, events, and orthologous gene blocks or orthoblocks have been described previously [25]. Briefly, the *reference taxon* is a strain in which the operon in question has been experimentally validated. Two genes are considered *neighboring genes* if they are 500 nucleotides or fewer apart and on the same strand. A *gene block* comprises no fewer than two such neighboring open reading frames. Organisms have *orthoblocks* when each has at least two neighboring genes that are homologous to genes in a gene block in the reference taxon’s genome. Using *Xanthomonas oryzae* pv. *oryzicola* BLS256 (*Xoc*) as a reference taxon, we retrieved the 10 genes in the GA operon (*cyp115, cyp112, cyp114, fd*_*GA*_, *sdr*_*GA*_, *cyp117, ggps, cps, ks*, and *idi*). From those 10 genes, we determined whether a query strain contains orthologous gene blocks. An *event* is a change in the gene block between any two species with homologous gene blocks. We identify three types of pairwise events between orthoblocks in different taxa: splits, deletions, and duplications. The event-based distance between any two orthoblocks is the sum of the minimized count of splits, duplications, and deletions.

#### Computational Reconstruction of the Gibberellin Operon Phylogeny

ROAGUE (Reconstruction of Ancestral Gene blocks Using Events) software was used to reconstruct ancestral gene blocks. ROAGUE accepts as input (1) a set of extant bacterial genomes, (2) a phylogenetic tree describing the relatedness between the set of species, and (3) a gold standard operon that has been experimentally validated from one species in the set of given genomes. ROAGUE finds the orthologs of the genes in the reference operons, then constructs the hypothesized ancestral gene blocks using a maximum parsimony algorithm, as previously described [33]. To assess the possibility of HGT among rhizobia species, phylogenetic trees were constructed using both a species marker gene (*rpoB*) and a concatenation of the protein sequences for genes in the GA operon. The topology for each of these reconstructions was then compared in order to find major incongruences between the two that may indicate HGT. For details see Figure 2 and the Supplementary Materials.

## ACKNOWLEDGMENTS

The authors thank Dr. Axel Schmidt and Prof. Jonathan Gershenzon (Max Planck Institute of Chemical Ecology) for use of their LC-MS/MS.

## FUNDING

This work was supported by grants to RJP from the NIH (GM076324) and USDA (NIFA-AFRI grant 2014-67013-21720), a postdoctoral fellowship to RN from the Deutsche Forschungsgemeinschaft (DFG) NA 1261/1-2, NSF ABI Development award 1458359 and NSF ABI Innovation award 1551363 to IF, a Discovery Grant and an Engage Grant, both from Natural Sciences and Engineering Research Council of Canada (NSERC) to TCC, and an Ontario Graduate Scholarship to AM.

## Notes

https://github.com/nguyenngochuy91/Ancestral-Blocks-Reconstruction

