## Supplementary Material for "Unraveling a tangled skein: Evolutionary analysis of the bacterial gibberellin biosynthetic operon"

### Supplementary Information

#### SUPPLEMENTARY METHODS

##### Biochemical characterization of *Re*IDS2 and *Et*IDI

*Re*IDS2 and *Et*IDI were cloned by amplification from genomic DNA of *Rhizobium etli* CE3 (a streptomycin-resistant derivative of *R. etli* CFN42)[1] or *Erwinia tracheiphila* PSU-1, respectively, with Q5 Hot Start High-Fidelity DNA polymerase (NEB) according to the product manual using gene specific primers (**Supplemental Table 3**) and 5 µL of the high GC-content enhancer for *Re*IDS2. The forward primer started with CACC to allow for directional cloning into pET101/D-TOPO(R) (Invitrogen), and each gene was sequenced verified following ligation into the vector.

For recombinant expression, pET101 containing either *Re*IDS or *Et*IDI were transformed into *E. coli* strain BL21 Star (Invitrogen). Starter cultures were inoculated into 10 mL NZY media (10 g $L^{-1}$ NaCl, 10 g $L^{-1}$ casein, 5 g $L^{-1}$ yeast extract, 1 g $L^{-1}$ anhydrous $MgSO_4$, pH 7.0) with 50 µg $mL^{-1}$ carbenicillin and grown at 200 rpm and 18 °C for 3 days. 5 ml from these starter cultures were used to inoculate 100 mL fresh NZY media containing 50 µg $mL^{-1}$ carbenicillin. After reaching an OD600 of 0.6, these were induced with 1 mM IPTG and grown under continuous shaking at 200 rpm at 18 °C for 24 hours. Cells were harvested by centrifugation at 5000 x *g* for 15 min. The cell pellet was resuspended in 5 mL MOPSO buffer (25 mM MOPSO, pH 7.2, 10 mM $MgCl_2$, 10% glycerol) with 20 mM imidazole, then lysed using an EmulsiFlex C-5 homogenizer (Avestin, Canada). The homogenized suspensions were centrifuged at 16,000 x *g* for 60 min. The supernatant was passed over 1 mL Ni-NTA agarose (Qiagen), which was then washed with 5 mL buffer containing 20 mM imidazole and then with an additional 5 mL of buffer with 50 mM imidazole. The recombinant 6xHis-tagged proteins were eluted with 2 mL buffer containing 250 mM imidazole.

IDS enzyme assays were carried out in triplicate with 10 µg of purified heterologously expressed protein in 300 µL of buffer and 50 µM IPP and 50 µM DMAPP as substrates. *Et*IDI assays were also performed in triplicate using a combined assay of 20 µg *Re*GGPS2 and 20 µg of *Et*IDI, with the addition of 10 µM flavin mononucleotide and 5 mM NADPH as described previously described[2] and either 100 µM IPP or DMAPP. Assays were incubated for 2 hrs at 30 °C, then flash frozen in liquid nitrogen and kept at -80°C until their analysis by LC-MS/MS, which was carried out as previously described.[3]

#### Analysis of GA operon GC content

GA operon sequences along with ~10 kilobases (kb) of genomic sequence flanking the operon on both sides were downloaded from NCBI/GenBank. Sequences were analyzed in Geneious® 10.2.3 (Biomatters, Ltd.), where GC content was determined for a sliding window size of 500 base pairs (bp).

#### Operon phylogenetic reconstruction

*Computational Reconstruction of the Gibberellin Operon Phylogeny*

The genome of *Xanthomonas oryzae* pv. oryzicola BLS256 (Xoc) was used as the gold standard, as it has a well-annotated genome, its GA operon contains full-length copies of all 10 archetypical GA operon genes (*cyp115*, *cyp112*, *cyp114*, *cyp117*, $fd_{\mathrm{GA}}$, $sdr_{\mathrm{GA}}$, *ggps*, *cps*, *ks*, and *idi*), and the majority of these have been biochemically characterized.[4] *Erwinia tracheiphila* PSU-1 was used as the phylogenetic outgroup, as this has been shown have the most distant GA operon.[5] ROAGUE was then applied to these two γ-proteobacteria (*E. tracheiphila* and Xoc) and α-proteobacteria rhizobia including *Azorhizobium*, *Bradyrhizobium*, *Mesorhizobium*, *Microvirga*, *Rhizobium*, and *Ensifer*/*Sinorhizobium* species. The full list of strains used is shown in the **Supplementary Table 4**.

Ancestral reconstruction was performed on both the species and operon sequence phylogenetic trees, in large part due to the previously observed incongruences between phylogenetic trees constructed with species markers (16S rRNA) and those found within the apparently horizontally transferred symbiotic module, as measured by analysis of *nifK* and GA operon genes.[6] Three different trees were used here to study the evolution of orthoblocks; a species tree, a concatenated operon tree, and a single operon gene tree using *cyp114*. The species tree *S*, was generated using the Neighbor-Joining algorithm with alignments of *rpoB* protein sequence as a species marker. For the operon tree *O*, the protein sequences of open reading frames (ORFs) of the orthoblock genes for each species were naively concatenated. Because many species lack a full length *cyp115* and *idi* gene, and due to the loss of *ggps* in a few species, only the following genes, which seem to be more uniformly conserved, were concatenated for this purpose: *cyp112*, *cyp114*, $fd_{GA}$, $sdr_{GA}$, *cyp117*, *cps*, and *ks*. A multiple sequence alignment of these concatenations was created, and the Neighbor-Joining algorithm was used to build the presented trees. Each leaf node *v* in *S*, and *O* contains orthologs to the genes found in the GA operon of the reference species (Xoc). For any two genes *a* and *b*, if the chromosomal distance is less than 500 bp, the genes will be written as *ab*. If the distance is greater than 500 bp, they are written with the separator character, thus written as *a*|*b*.

Due to the large number (118) and redundancy (both phylogenetically and in operon structure) of the rhizobial strains, the size of the tree was reduced by using the Phylogenetic Diversity Analyzer software.[7] To facilitate analysis and presentation, the number of species was reduced to 64, which was still representative of the overall diversity, and enables ready visualization. Additionally, this software only keeps species that are representative enough to give distinct branches on the phylogenetic tree (i.e., reflecting appreciable distance between species). Accordingly, this approach also eliminated redundancy that would otherwise have confounded this analysis. The full operon tree, full species tree, partial operon tree, and partial species tree are referred to as *FO*, *FS*, *PO*, *PS* trees, respectively.

*Treatment of gene fusion events*

Currently, ROAGUE does not account for gene fusion. Given the presence of several GA operons wherein the $fd_{GA}$ gene is fused in frame with either the $sdr_{GA}$ gene or the *cyp114* gene, it was necessary to assess these manually. This was accomplished by rechecking the gene block to determine if the $fd_{GA}$ gene was missing, as the fusion of this gene in frame with either $sdr_{GA}$ or *cyp114* would result in it not being recognized as a unique ORF by the ROAGUE software. Then, the BLAST results were queried and potential fusions found by checking the start and end of the alignments for the subject genome. In addition, this approach was used to distinguish pseudo-genes or fragments of *cyp115* from the full-length gene, if present, by accounting for nucleotide identity and alignment length.

**SUPPLEMENTARY FIGURES**

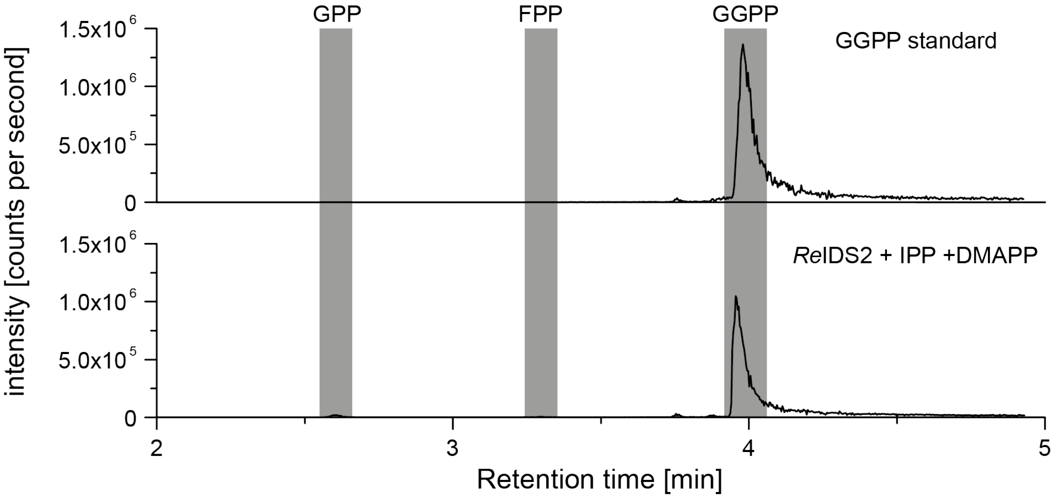


**Supplementary Figure 1. *In vitro* characterization of ReIDS2.** LC-MS/MS chromatograms for the *in vitro* enzyme assay of *Re*IDS2 with IPP and DMAPP as substrates (bottom panel) in comparison to an authentic GGPP standard (top panel). Note that trace amounts of geranyl diphosphate (GDP) and farnesyl diphosphate (FDP) could be detected as products of the enzyme assay. Intensity of peaks is measured in ion counts per second.

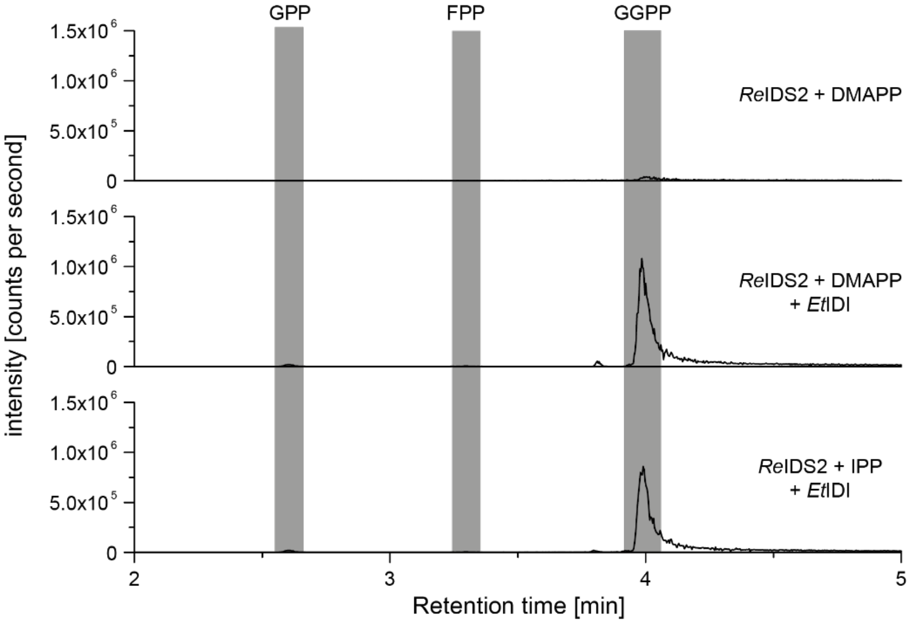


**Supplementary Figure 2. *In vitro* characterization of *Et*IDI.** LC-MS/MS chromatograms for the *in vitro* enzyme assay of *Re*IDS2 with DMAPP as substrate (top panel); *Re*IDS2 and *Et*IDI combination assay with DMAPP as substrate (middle panel); *Re*IDS2 and *Et*IDI combination assay with IPP as substrate (bottom panel). Intensity of peaks is measured in ion counts per second.

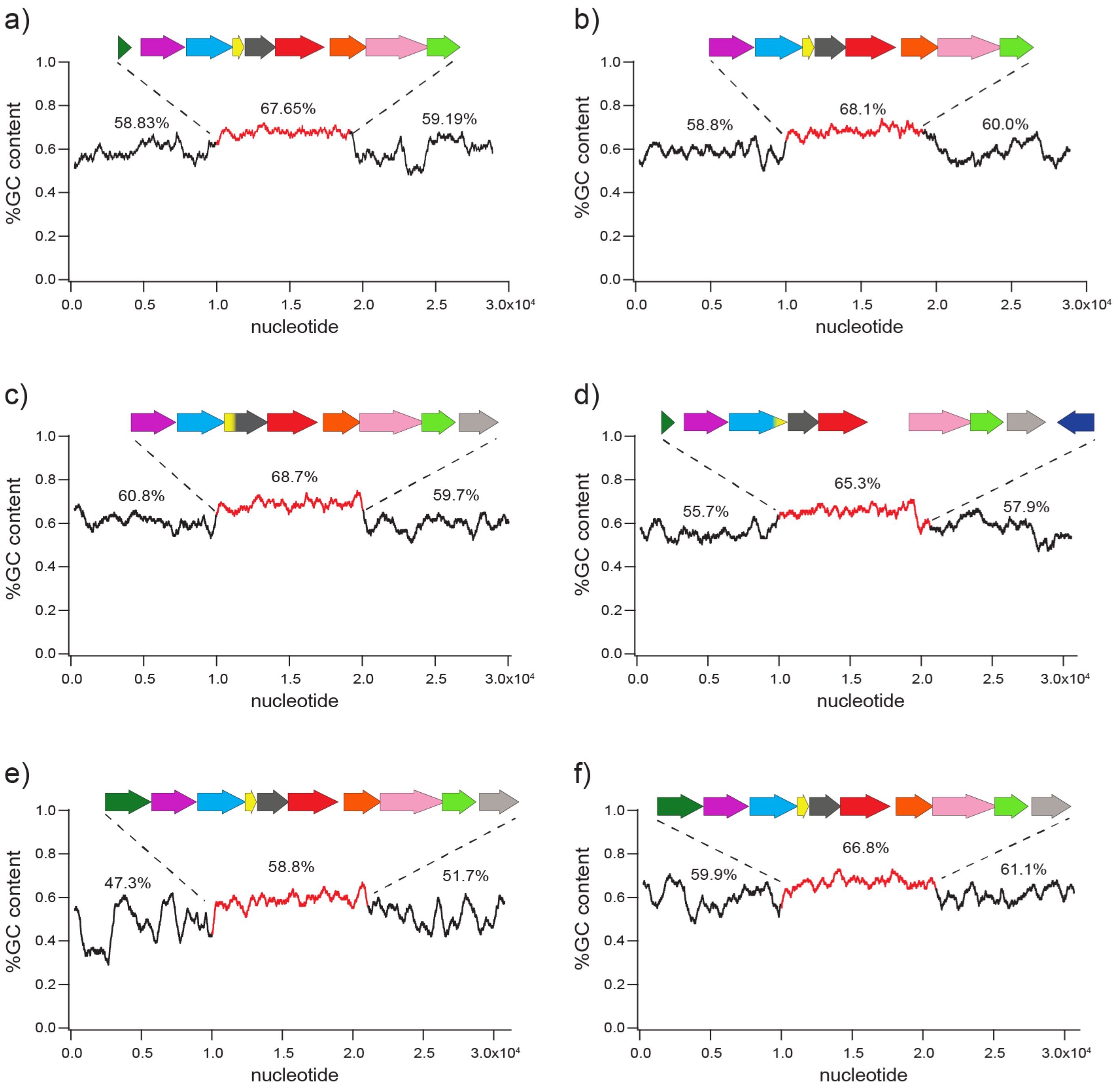


**Supplemental Figure 3**. **GC content analysis of the GA operon in a range of bacterial lineages**. The average percent GC content within 500 base pair windows is shown for the GA operon (red) and flanking 10 kilobase regions for the following species: a) *B. japonicum* USDA 110, b) *S. fredii* NGR234, c) *M. loti* MAFF303099, d) *R. etli* CFN 42, e) *E. tracheiphila* PSU-1, and f) *X. oryzae* pv. *oryzicola* BLS256. Shown above each region is the average percent GC content.

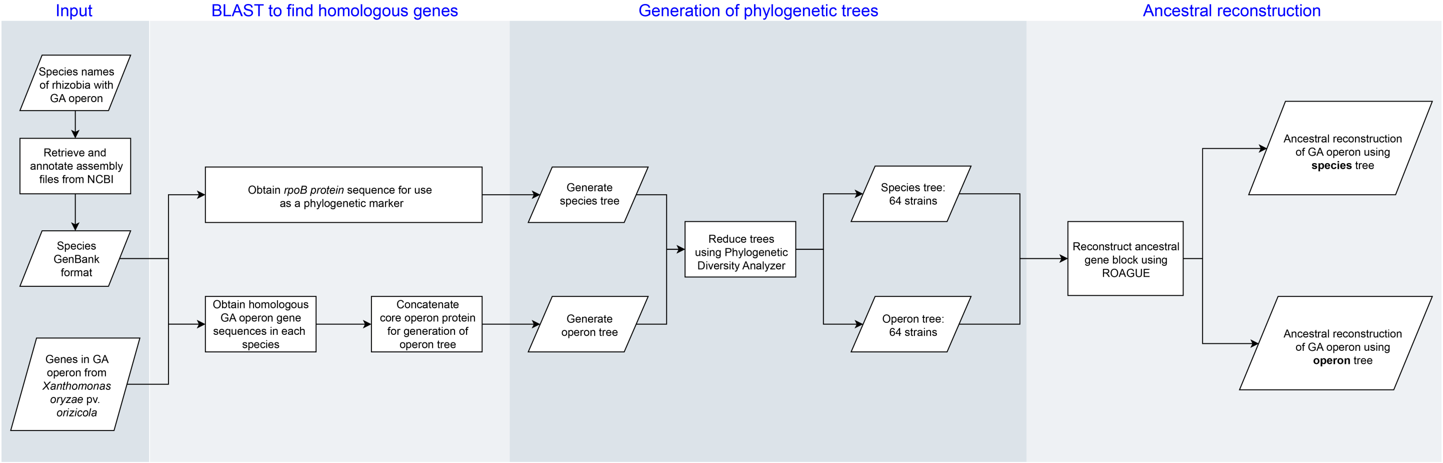


**Supplementary Figure 4. Method pipeline the ROAGUE-generated ancestral operon reconstructions.**

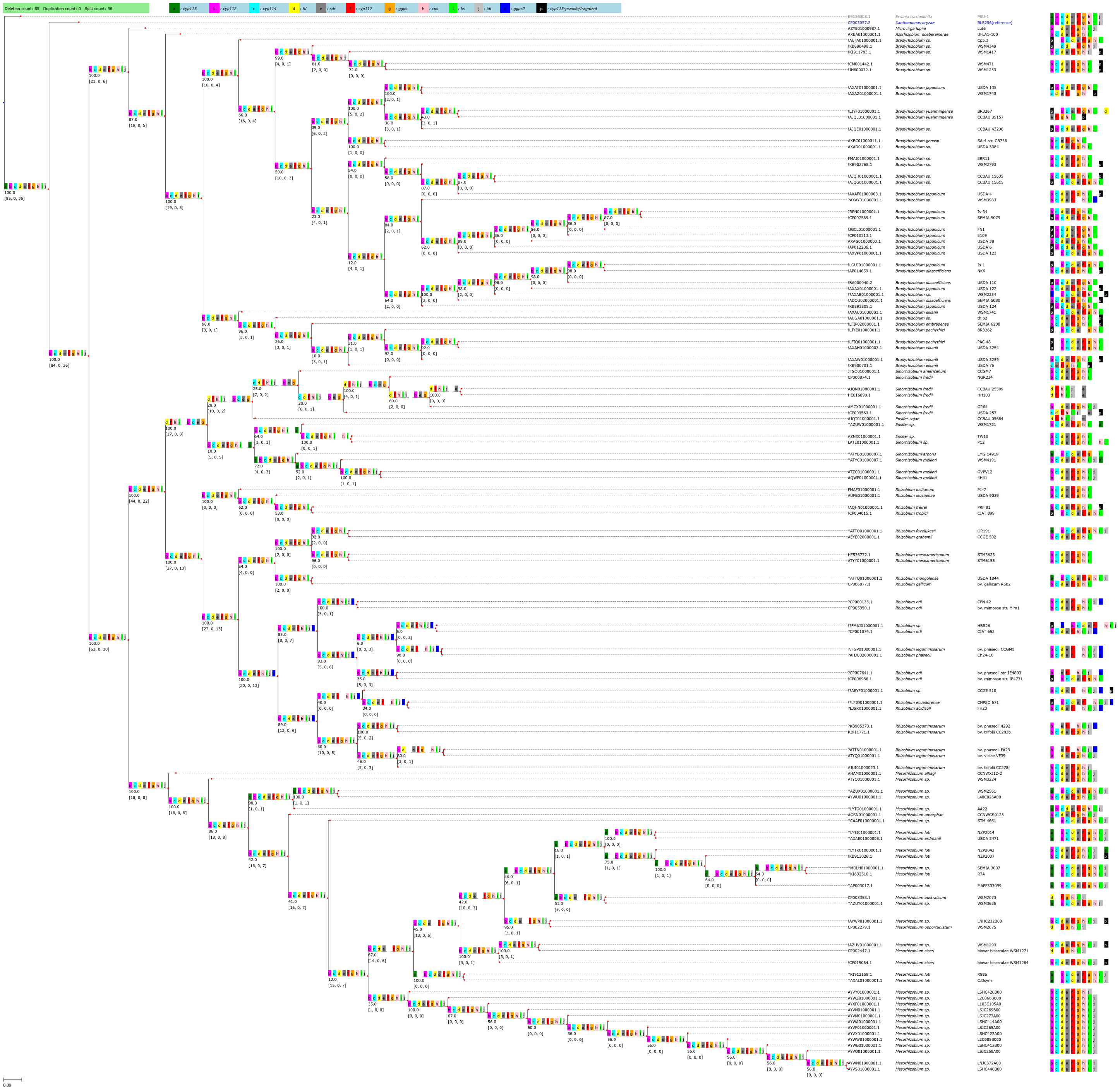


**Supplementary Figure 5**. **Ancestral reconstruction of the GA biosynthetic operon using *rpoB* for phylogenetic analysis.** A phylogenetic tree was constructed with alignments of *rpoB* protein sequences from 118 α-rhizobia species and two γ-proteobacteria using the Neighbor-Joining method as a measure of distance between species. ROAGUE was then applied to create the ancestral operon reconstruction. The lower-case letters in each tree node represent the genes in the orthoblock (e.g. "a" represents "*cyp115*"), with each gene additionally indicated by a unique color (see legend at top of figure). A blank space between genes designates a split ≥500bp between the genes to either side of the blank space. The green bar on the top left of the figure displays the total number of events that occur in this reconstruction. For each inner node *u*, the floating number (e.g. 98.0) represents the bootstrap value of the tree. The numbers in the brackets indicate the cumulative count of events going from the leaf nodes to node *u* in the following order: [deletions, duplications, splits]. Each leaf node is accompanied with symbols (*, ?, !), the genomic accession number, the species/strain name, and the gene block for that strain. An asterisk (*) indicates the gene block contains full length *cyp115* (gene "a"); an exclamation (!) indicates that the gene block contains a truncation/fragment of *cyp115*, and a question mark (?) indicates the gene block contains *ggps2* (gene "k"). The reference strain, *Xanthomonas oryzae* pv. *oryzicola* BLS256, is in blue, and the outgroup strain, *Erwinia tracheiphila* PSU-1, is in gray. These naming and color conventions persist through this study.

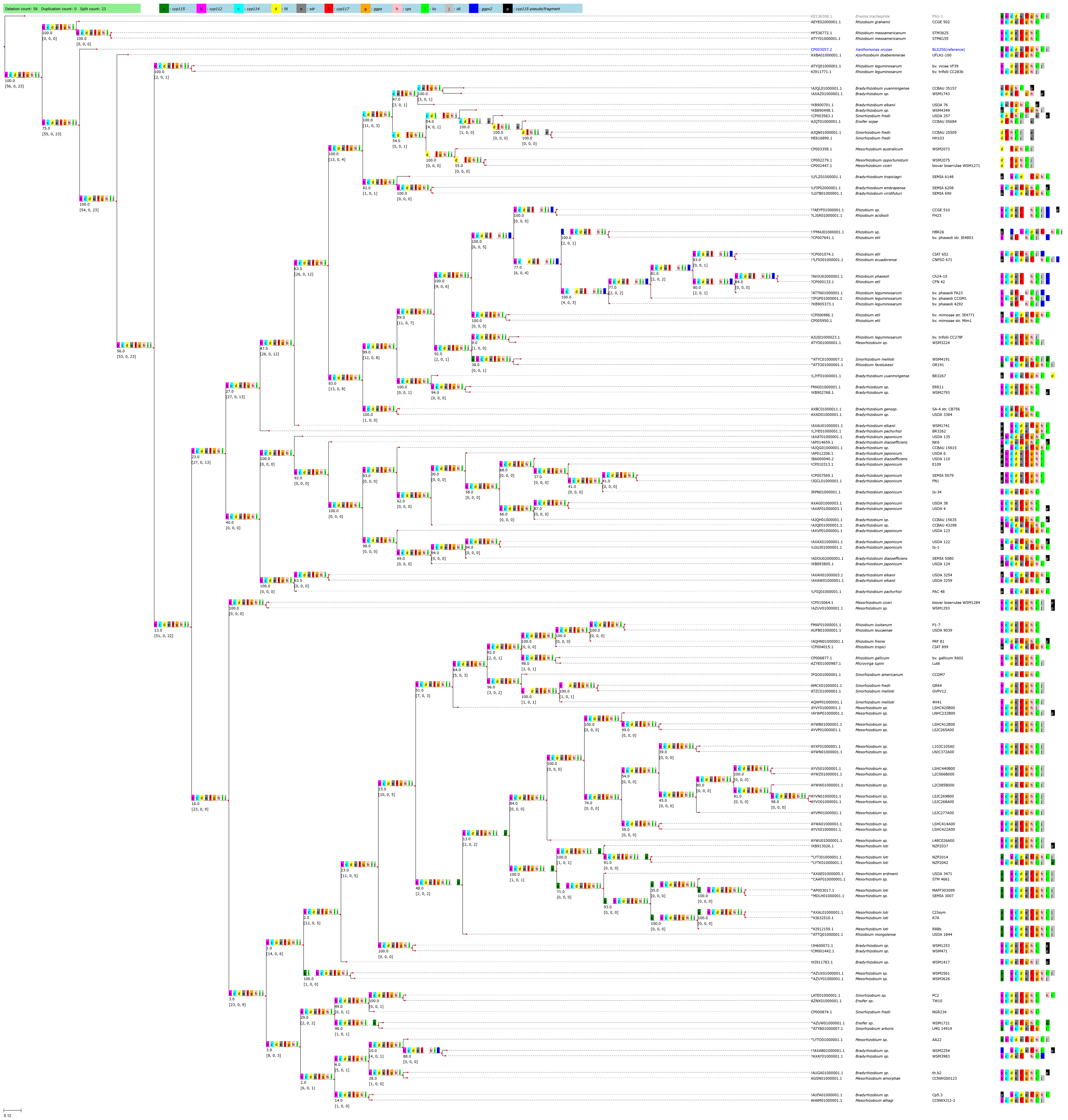

**Supplementary Figure 6**. **Ancestral reconstruction of the GA biosynthetic operon using the concatenated operon for phylogenetic analysis.** A phylogenetic tree was constructed with alignments of concatenated proteins from core GA operon genes (*cyp112-cyp114-fd-sdr-cyp117-cps-ks*) from 118 α-rhizobia species and two γ-proteobacteria using the Neighbor Joining method. ROAGUE was then applied to create the ancestral operon reconstruction. ROAGUE was then applied to create the ancestral operon reconstruction. All annotations are described previously in **Supplementary Figure 4**.

**Supplementary Table 1. Strains containing a CYP114-$Fd_{GA}$ gene fusion.**

| |
|---|
| *Bradyrhizobium* sp. ERR11 |
| *Bradyrhizobium yuanmingense* BR3267 |
| *Rhizobium etli* CIAT 652 |
| *Rhizobium etli* CNPAF512 |
| *Rhizobium etli* bv. *phaseoli* IE4803 |
| *Rhizobium phaseoli* Ch24-10 |
| *Rhizobium etli* CFN 42 |
| *Rhizobium leguminosarum* bv. *phaseoli* CCGM1 |
| *Rhizobium acidisoli* FH23 |
| *Rhizobium* sp. HBR26 |
| *Mesorhizobium* sp. WSM3224 |
| *Rhizobium etli* bv. *phaseoli* str. IE4803 |
| *Rhizobium leguminosarum* bv. *trifolii* CC278f |
| *Rhizobium leguminosarum* bv. *phaseoli* FA23 |
| *Rhizobium etli* bv. *mimosae* str. Mim1 |
| *Sinorhizobium meliloti* WSM4191 |

**Supplementary Table 2. Strains containing a $Fd_{GA}$-$SDR_{GA}$ gene fusion.**

| |
|---|
| *Mesorhizobium* sp. LSHC414A00 |
| *Mesorhizobium loti* MAFF303099 |
| *Mesorhizobium loti* CJ3sym |
| *Mesorhizobium loti* R7A |
| *Rhizobium mongolense* USDA 1844 |
| *Mesorhizobium* sp. LSJC269B00 |
| *Mesorhizobium* sp. LSJC277A00 |
| *Mesorhizobium* sp. LSJC265A00 |

| |
|---|
| *Mesorhizobium* sp. L2C066B000 |
| *Mesorhizobium* sp. L2C085B000 |
| *Bradyrhizobium* sp. WSM1253 |
| *Mesorhizobium loti* NZP2042 |
| *Mesorhizobium* sp. LSHC440B00 |
| *Mesorhizobium* sp. SEMIA_3007 |
| *Mesorhizobium* sp. L48C026A00 |
| *Mesorhizobium* sp. LNHC232B00 |
| *Bradyrhizobium* sp. WSM471 |
| *Bradyrhizobium japonicum* USDA 135 |
| *Mesorhizobium* sp. LSJC268A00 |
| *Mesorhizobium* sp. STM 4661 |
| *Mesorhizobium loti* NZP2037 |
| *Mesorhizobium loti* R88b |
| *Mesorhizobium loti* NZP2014 |
| *Mesorhizobium* sp. LSHC420B00 |
| *Mesorhizobium* sp. LNJC372A00 |
| *Mesorhizobium* sp. LSHC412B00 |
| *Mesorhizobium erdmanii* USDA 3471 |
| *Mesorhizobium* sp. L103C105A0 |
| *Mesorhizobium* sp. LSHC422A00 |

**Supplemental Table 3. List of primers used in this study.**

| Primer name | Primer Sequence |
|---|---|
| *E. tracheiphila* IDI pET forward | CACC ATG AAA CCG AGT GAT ACT CTA AGT CAA CGC AAG G |
| *E. tracheiphila* IDI reverse | CAC GGG GAT AAT GCC GGG AGG |
| *R. etli* IDS2 pET forward | CACC ATG ATT TCG AAT CAC CAA GCC GAC GTG G |
| *R. etli* IDS2 pET reverse | CAA AGG CGC GCA ATC CAG CG |

**Supplementary Table 4. List of strains used within the ancestral reconstruction analyses.** Shown for each strain is the corresponding GenBank genome accession, the legume host from which it was isolated, and the nodule type (D = determinate, I = indeterminate, * = neither). Note that the legume host and nodule type are irrelevant for the gammaproteobacteria plant pathogens analyzed in these reconstructions.

| GenBank genome accession | strain | legume host | nodule type |
|---|---|---|---|
| AXBA01000001.1 | *Azorhizobium doebereinerae* UFLA1-100 | *Sesbania virgata* | * |
| AP014659.1 | *Bradyrhizobium diazoefficiens* NK6 | *Glycine max* | D |
| ADOU02000001.1 | *Bradyrhizobium diazoefficiens* SEMIA 5080 | *Glycine max* | D |
| BA000040.2 | *Bradyrhizobium diazoefficiens* USDA 110 | *Glycine max* | D |
| AXAH01000003.1 | *Bradyrhizobium elkanii* USDA 3254 | *Phaseolus acutifolius* | D |
| AXAW01000001.1 | *Bradyrhizobium elkanii* USDA 3259 | *Phaseolus lunatus* | D |
| KB900701.1 | *Bradyrhizobium elkanii* USDA 76 | *Glycine max* | D |
| AXAU01000001.1 | *Bradyrhizobium elkanii* WSM1741 | *Rhynchosia minima* | D |
| LFIP02000001.1 | *Bradyrhizobium embrapense* SEMIA 6208 | *Desmodium heterocarpon* | D |
| AXBC01000011.1 | *Bradyrhizobium* genosp. SA-4 str. CB756 | *Macrotyloma africanum* | D |
| CP010313.1 | *Bradyrhizobium japonicum* E109 | *Glycine max* | D |
| JGCL01000001.1 | *Bradyrhizobium japonicum* FN1 | *Glycine max* (soil isolate) | D |
| LGUJ01000001.1 | *Bradyrhizobium japonicum* Is-1 | *Glycine max* | D |
| JRPN01000001.1 | *Bradyrhizobium japonicum* Is-34 | *Glycine max* | D |
| CP007569.1 | *Bradyrhizobium japonicum* SEMIA 5079 | *Glycine max* | D |
| AXAX01000001.1 | *Bradyrhizobium japonicum* USDA 122 | *Glycine max* | D |
| AXVP01000001.1 | *Bradyrhizobium japonicum* USDA 123 | *Glycine max* | D |
| KB893805.1 | *Bradyrhizobium japonicum* USDA 124 | *Glycine max* | D |
| AXAT01000001.1 | *Bradyrhizobium japonicum* USDA 135 | *Glycine max* | D |
| AXAG01000003.1 | *Bradyrhizobium japonicum* USDA 38 | *Glycine max* | D |
| AXAF01000003.1 | *Bradyrhizobium japonicum* USDA 4 | *Glycine max* | D |
| AP012206.1 | *Bradyrhizobium japonicum* USDA 6 | *Glycine max* | D |
| LJYE01000001.1 | *Bradyrhizobium pachyrhizi* BR3262 | *Vigna unguiculata* | D |
| LFIQ01000001.1 | *Bradyrhizobium pachyrhizi* PAC 48 | *Pachyrhizus erosus* | D |
| AJQG01000001.1 | *Bradyrhizobium* sp. CCBAU 15615 | *Glycine max* | D |
| AJQH01000001.1 | *Bradyrhizobium* sp. CCBAU 15635 | *Glycine max* | D |
| AJQE01000001.1 | *Bradyrhizobium* sp. CCBAU 43298 | *Glycine max* | D |
| AUFA01000001.1 | *Bradyrhizobium* sp. Cp5.3 | *Centrosema pubescens* | D |
| FMAI01000001.1 | *Bradyrhizobium* sp. ERR11 | *Erythrina brucei* | D |
| AUGA01000001.1 | *Bradyrhizobium* sp. th.b2 | *Amphicarpaea bracteata* | D |
| AXAD01000001.1 | *Bradyrhizobium* sp. USDA 3384 | *Kennedia coccinea* | D |
| JH600072.1 | *Bradyrhizobium* sp. WSM1253 | *Ornithopus compressus* | D |
| KI911783.1 | *Bradyrhizobium* sp. WSM1417 | *Lupinus sp.* | I |
| AXAZ01000001.1 | *Bradyrhizobium* sp. WSM1743 | *Indigofera* sp. | I |
| AXAB01000001.1 | *Bradyrhizobium* sp. WSM2254 | *Acacia dealbata* | I |
| KB902768.1 | *Bradyrhizobium* sp. WSM2793 | *Rhynchosia totta* | D |
| AXAY01000001.1 | *Bradyrhizobium* sp. WSM3983 | *Kennedia coccinea* | D |
| KB890498.1 | *Bradyrhizobium* sp. WSM4349 | *Syrmatium glabrum (Lotus scoparius)* | D* |

| | | | |
|---|---|---|---|
| CM001442.1 | *Bradyrhizobium* sp. WSM471 | *Ornithopus pinnatus* | D |
| LJYF01000001.1 | *Bradyrhizobium yuanmingense* BR3267 | *Kennedia coccinea* | D |
| AJQL01000001.1 | *Bradyrhizobium yuanmingense* CCBAU 35157 | *Glycine max* | D |
| AJQT01000001.1 | *Ensifer sojae* CCBAU 5684 | *Glycine max* | D |
| AZNX01000001.1 | *Ensifer* sp. TW10 | *Tephrosia purpurea* | I |
| AZUW01000001.1 | *Ensifer* sp. WSM1721 | *Indigofera sp.* | I |
| AHAM01000001.1 | *Mesorhizobium alhagi* CCNWXJ12-2 | *Alhagi sparsifolia* | I |
| AGSN01000001.1 | *Mesorhizobium amorphae* CCNWGS0123 | *Robinia pseudoacacia* | I |
| CP003358.1 | *Mesorhizobium australicum* WSM2073 | *Biserrula pelecinus* | I |
| CP002447.1 | *Mesorhizobium ciceri* bv. *biserrulae* WSM1271 | *Biserrula pelecinus* | I |
| CP015064.1 | *Mesorhizobium ciceri* bv. *biserrulae* WSM1284 | *Biserrula pelecinus* | I |
| AXAE01000005.1 | *Mesorhizobium erdmanii* USDA 3471 | *Lotus corniculatus* | D |
| AXAL01000001.1 | *Mesorhizobium loti* CJ3sym | *Lotus corniculatus* | D |
| AP003017.1 | *Mesorhizobium loti* MAFF303099 | *Lotus pedunculatus* | D |
| LYTJ01000001.1 | *Mesorhizobium loti* NZP2014 | *Lotus sp.* | D |
| KB913026.1 | *Mesorhizobium loti* NZP2037 | *Lotus divaricatus* | D |
| LYTK01000001.1 | *Mesorhizobium loti* NZP2042 | *Lotus sp.* | D |
| KI632510.1 | *Mesorhizobium loti* R7A | *Lotus corniculatus* | D |
| KI912159.1 | *Mesorhizobium loti* R88b | *Lotus corniculatus* | D |
| CP002279.1 | *Mesorhizobium opportunistum* WSM2075 | *Biserrula pelecinus* | I |
| LYTO01000001.1 | *Mesorhizobium* sp. AA22 | *Biserrula pelecinus L* | I |
| AYXF01000001.1 | *Mesorhizobium* sp. L103C105A0 | *Acmispon wrangelianus* | D |
| AYWZ01000001.1 | *Mesorhizobium* sp. L2C066B000 | *Acmispon wrangelianus* | D |
| AYWW01000001.1 | *Mesorhizobium* sp. L2C085B000 | *Acmispon wrangelianus* | D |
| AYWU01000001.1 | *Mesorhizobium* sp. L48C026A00 | *Acmispon wrangelianus* | D |
| AYWP01000001.1 | *Mesorhizobium* sp. LNHC232B00 | *Acmispon wrangelianus* | D |
| AYWN01000001.1 | *Mesorhizobium* sp. LNJC372A00 | *Acmispon wrangelianus* | D |
| AYWB01000001.1 | *Mesorhizobium* sp. LSHC412B00 | *Acmispon wrangelianus* | D |
| AYWA01000001.1 | *Mesorhizobium* sp. LSHC414A00 | *Acmispon wrangelianus* | D |
| AYVY01000001.1 | *Mesorhizobium* sp. LSHC420B00 | *Acmispon wrangelianus* | D |
| AYVX01000001.1 | *Mesorhizobium* sp. LSHC422A00 | *Acmispon wrangelianus* | D |
| AYVS01000001.1 | *Mesorhizobium* sp. LSHC440B00 | *Acmispon wrangelianus* | D |
| AYVP01000001.1 | *Mesorhizobium* sp. LSJC265A00 | *Acmispon wrangelianus* | D |
| AYVO01000001.1 | *Mesorhizobium* sp. LSJC268A00 | *Acmispon wrangelianus* | D |
| AYVN01000001.1 | *Mesorhizobium* sp. LSJC269B00 | *Acmispon wrangelianus* | D |
| AYVM01000001.1 | *Mesorhizobium* sp. LSJC277A00 | *Acmispon wrangelianus* | D |
| MDLH01000001.1 | *Mesorhizobium* sp. SEMIA 3007 | *Pisum sativum* | I |
| CAAF010000001.1 | *Mesorhizobium* sp. STM 4661 | *Anthyllis vulneraria* | D |
| AZUV01000001.1 | *Mesorhizobium* sp. WSM1293 | *Lotus sp.* | D |
| AZUX01000001.1 | *Mesorhizobium* sp. WSM2561 | *Lessertia diffusa* | I |
| ATYO01000001.1 | *Mesorhizobium* sp. WSM3224 | *Otholobium candicans* | D |
| AZUY01000001.1 | *Mesorhizobium* sp. WSM3626 | *Lessertia diffusa* | I |
| AZYE01000987.1 | *Microvirga lupini* Lut6 | *Lupinus texensis* | I |
| LJSR01000001.1 | *Rhizobium acidisoli* FH23 | *Phaseolus vulgaris* | D |
| LFIO01000001.1 | *Rhizobium ecuadorense* CNPSO 671 | *Phaseolus vulgaris* | D |
| CP006986.1 | *Rhizobium etli* bv. *mimosae* str. IE4771 | *Phaseolus vulgaris* | D |
| CP005950.1 | *Rhizobium etli* bv. *mimosae* str. Mim1 | *Mimosa affinis* | I |
| CP007641.1 | *Rhizobium etli* bv. *phaseoli* str. IE4803 | *Phaseolus vulgaris* | D |

| | | | |
|---|---|---|---|
| CP000133.1 | *Rhizobium etli* CFN 42 | *Phaseolus vulgaris* | D |
| CP001074.1 | *Rhizobium etli* CIAT 652 | *Phaseolus vulgaris* | D |
| ATTO01000001.1 | *Rhizobium favelukesii* OR191 | *Medicago sativa* | I |
| AQHN01000001.1 | *Rhizobium freirei* PRF 81 | *Phaseolus vulgaris* | D |
| CP006877.1 | *Rhizobium gallicum* bv. *gallicum* R602 | *Phaseolus vulgaris* | D |
| AEYE02000001.1 | *Rhizobium grahamii* CCGE 502 | *Dalea leporin* | I |
| KB905373.1 | *Rhizobium leguminosarum* bv. *phaseoli* 4292 | *Phaseolus vulgaris* | D |
| JFGP01000001.1 | *Rhizobium leguminosarum* bv. *phaseoli* CCGM1 | *Phaseolus vulgaris* | D |
| ATTN01000001.1 | *Rhizobium leguminosarum* bv. *phaseoli* FA23 | *Phaseolus vulgaris* | D |
| AJUI01000023.1 | *Rhizobium leguminosarum* bv. *trifolii* CC278f | *Trifolium nanum* | I |
| KI911771.1 | *Rhizobium leguminosarum* bv. *trifolii* CC283b | *Trifolium ambiguum* | I |
| ATYQ01000001.1 | *Rhizobium leguminosarum* bv. *viciae* VF39 | *Vicia faba* | I |
| AUFB01000001.1 | *Rhizobium leucaenae* USDA 9039 | *Phaseolus vulgaris* | D |
| FMAF01000001.1 | *Rhizobium lusitanum* P1-7 | *Phaseolus vulgaris* | D |
| HF536772.1 | *Rhizobium mesoamericanum* STM3625 | *Mimosa pudica* | I |
| ATYY01000001.1 | *Rhizobium mesoamericanum* STM6155 | *Mimosa pudica* | I |
| ATTQ01000001.1 | *Rhizobium mongolense* USDA 1844 | *Medicago ruthenica* | I |
| AHJU02000001.1 | *Rhizobium phaseoli* Ch24-10 | *Phaseolus vulgaris* (maize stem isolate) | D |
| AEYF01000001.1 | *Rhizobium* sp. CCGE 510 | *Phaseolus albescens* | D |
| FMAJ01000001.1 | *Rhizobium* sp. HBR26 | *Phaseolus vulgaris* | D |
| CP004015.1 | *Rhizobium tropici* CIAT 899 | *Phaseolus vulgaris* | D |
| JFGO01000001.1 | *Sinorhizobium americanum* CCGM7 | *Phaseolus vulgaris* | D |
| ATYB01000007.1 | *Sinorhizobium arboris* LMG 14919 | *Prosopis chilensis* | I |
| AJQN01000001.1 | *Sinorhizobium fredii* CCBAU 25509 | *Glycine max* | D |
| AMCX01000001.1 | *Sinorhizobium fredii* GR64 | *Phaseolus vulgaris* | D |
| HE616890.1 | *Sinorhizobium fredii* HH103 | *Glycine max* (soil isolate) | D |
| CP000874.1 | *Sinorhizobium fredii* NGR234 | *Lablab purpureus* | D |
| CP003563.1 | *Sinorhizobium fredii* USDA 257 | *Glycine max* | D |
| AQWP01000001.1 | *Sinorhizobium meliloti* 4H41 | *Phaseolus vulgaris* | D |
| ATZC01000001.1 | *Sinorhizobium meliloti* GVPV12 | *Phaseolus vulgaris* | D |
| ATYC01000007.1 | *Sinorhizobium meliloti* WSM4191 | *Melilotus siculus* | I |
| LATE01000001.1 | *Sinorhizobium* sp. PC2 | *Prosopis cineraria* | I |
| CP003057.2 | *Xanthomonas oryzae* pv. *oryzicola* BLS256 | - | - |
| KE136308.1 | *Erwinia tracheiphila* PSU-1 | - | - |
